# *DROSOPHILA* MTOR COMPLEX 2 PRESERVES MITOCHONDRIAL AND CARDIAC FUNCTION UNDER HIGH FAT DIET TREATMENT

**DOI:** 10.1101/2021.01.27.428443

**Authors:** Kai Chang, Guillermo A. Requejo Figueroa, Hua Bai

## Abstract

High fat diet (HFD)-associated lipotoxicity is one of the major causes of cardiovascular diseases. The mechanistic target of rapamycin (mTOR) pathway, especially mTOR complex 1 (mTORC1), has been previously implicated in HFD-induced heart dysfunction. In the present study, we find that unlike mTORC1, mTOR complex 2 (mTORC2) protects hearts from HFD-induced cardiomyopathy and mitochondrial dysfunction in *Drosophila*. We show that HFD feeding induces contractile dysfunction along with altered mitochondrial morphology and function. Upon HFD feeding, the mitochondria of cardiomyocytes exhibit fragmentation, loss of membrane potential, and calcium overload. Interestingly, HFD feeding also reduces the activity of cardiac mTORC2. In line with this finding, the flies with cardiac-specific knockdown of *rictor*, the key subunit of mTORC2, show cardiac and mitochondrial dysfunction similar to what is observed in HFD-fed wild-type flies. Conversely, cardiac-specific activation of mTORC2 by overexpressing *rictor* attenuates HFD-induced mitochondrial and cardiac dysfunction. Thus, our findings suggest that mTORC2 is a cardioprotective factor and regulates mitochondrial homeostasis upon HFD feeding.

## Introduction

Obesity has grown to pandemic levels with nearly three folds increases since 1975 (Blüher 2019). Increasing evidence suggests that obesity and its associated metabolic disorders caused by excessive fat intake increase the risk of developing secondary diseases such as type-2 diabetes and cardiovascular diseases (Birse and Bodmer 2011). Obese people and type-2 diabetic patients exhibit several cardiac dysfunctions including ventricular remodeling, diastolic/systolic dysfunction, decreased fractional shortening, and prolonged QT intervals (Christoffersen, Bollano et al. 2003, Birse, Choi et al. 2010, Birse and Bodmer 2011, Zhang and Ren 2011). Cardiomyocytes require a constant supply of energy in the form of adenosine triphosphate (ATP) to support its contractile function. Under normal condition, most ATP in cardiomyocytes is generated through β-oxidation of free fatty acids (FFAs). During the development of obesity due to high caloric intake, the availability of FFAs is increased in the heart, which in turn promotes fatty acid oxidation that eventually leads to contractile dysfunction (Lopaschuk, Folmes et al. 2007, Birse, Choi et al. 2010).

Mitochondrial dysfunction contributes significantly to the progression of cardiomyopathy under nutrient overload (Boudina, Sena et al. 2007, Lopaschuk, Folmes et al. 2007). Mitochondria are highly dynamic organelles and their morphology and function are often altered upon high fat diet (HFD) treatment, such as fragmented mitochondria, decreased complex I activity, and induction of mitophagy. In contrast, other studies reported that mitochondrial function was unaffected or even increased after feeding[16], [17]. Therefore, it is necessary to define the precise mitochondrial responses to HFD so that we could better understand the mechanisms underlying HFD-induced mitochondrial changes, especially in the heart.

The mechanistic target of rapamycin (mTOR) pathway is a highly conserved nutrient-sensing pathway that functions through two structurally and functionally distinct complexes, mTOR complex 1 (mTORC1) and mTOR complex 2 (mTORC2), to regulate a wide range of cellular function including protein synthesis, ribosomal and mitochondrial biogenesis, autophagy, and metabolism[18], [19]. Abundant evidence suggests that obesity and nutrient overload induce a hyper-activation of mTOR activity in multiple tissues, contributing to the development of type-2 diabetes and insulin resistance[20]. Recently, a study in *Drosophila* showed that mTOR signaling also plays a central role in HFD-induced heart dysfunction. Reducing insulin-mTOR pathway activity prevents HFD-induced triglyceride levels and cardiac abnormalities[5]. Additionally, increasing AMPK/mTOR pathway activity has also been observed in rats fed with HFD to mediate vascular dysfunction and remodeling [21]. However, the above studies did not differentiate the effects coming from two mTOR complexes. To our knowledge, most of the genetic manipulation that induces mTOR activity is achieved by activating mTORC1 activity.

Compared with mTORC1, the upstream signals and downstream substrates of mTORC2 are less known. Recently studies suggest that mTORC2 might also play a role in HFD-induced obesity and insulin resistance via an unknown mechanism[20], [22]–[24]. For example, HFD significantly decreases the protein levels of mTORC2 and pAKT, which is opposite to the protein levels of mTORC1[22]. Mice with mTORC2 deficiency display glucose tolerance that is generally observed in HFD treatment[25]. A recent study in neurons suggests that mTORC2 might also affect how rewarding high fat foods are[23]. Interestingly, a recent study has shown that mTORC2 localizes to mitochondrial-associated endoplasmic reticulum (ER)-membranes (MAMs) to regulate mitochondrial physiology[26]. As its name indicated, MAMs represent a region where ER makes contact with mitochondria. MAMs are involved in importing the lipid and calcium from the ER to mitochondria and regulating mitochondrial dynamics and metabolism[27]. Moreover, this crosstalk between mitochondria and ER is a prerequisite for healthy cardiac function[28], [29]. Therefore, it is likely that mTORC2 might regulate HFD-induced obesity and cardiac dysfunction via mediating mitochondrial physiology at MAMs. Collectively, mTORC2 seems to play a different, or even an opposite role to mTORC1 in the regulation of HFD-induced obesity, which requires further investigation.

*Drosophila melanogaster* has recently emerged as a suitable model to investigate the genetic mechanisms underlying HFD-induced obesity and cardiac dysfunction[30]–[34]. *Drosophil*a fed a HFD exhibit increased triglyceride fat, deregulation of insulin-mTOR signaling, insulin resistance, oxidative stress, metabolic inflexibility, and cardiac dysfunction. A recent study in *Drosophila* skeletal muscles showed that mitochondrial respiration is also affected by HFD treatment even though measurements on other mitochondrial physiology such as mitochondrial morphology and membrane potential are still lacking[17]. In this study, we used *Drosophila* as our model to investigate mitochondrial responses under HFD, especially in the heart, and the role of mTORC2 in regulating these responses. Our results indicated that mTORC2 could provide cardio-protection in response to HFD.

## Results

To better understand how HFD affects heart function in *Drosophila*, we first investigate the cardiac mitochondrial physiology and cardiac contractile function after feeding a HFD. Specifically, we fed *Drosophila* either a standard diet (SD) or a HFD (SD supplemented with 20% (w/v) coconut oil) for five days before the measurement. Several studies suggest that HFD induces a shift toward mitochondrial fission (Chen, Li, Zhang, Zhu, & Gao, 2018;Jheng et al., 2012; Leduc-Gaudet et al., 2018). Consistently, we found that the cardiac mitochondria became fragmented upon five days HFD treatment, indicated by a significantly increased number of smaller mitochondria in the HFD-treated heart (Figure 1A, B). Mitochondrial membrane potential (ΔΨm), as an essential component in oxidative phosphorylation, is a crucial indicator of mitochondrial activity, especially for cells with high ATP demand, such as cardiomyocytes, where ATP turnover has greater control over mitochondrial respiration and mitochondrial membrane potential. Therefore, we measured the mitochondrial membrane potential by tetramethylrhodamine ethyl ester (TMRE), a cell-permeable and cationic red-orange dye, in *Drosophila* heart fed with a HFD as well. We found that five days of HFD feeding significantly decreased ΔΨm in *Drosophila* heart than SD, suggesting that HFD impairs the cardiac mitochondrial respiration (Figure 1C, D).

**Table 1.** Time-dependent HFD-induced mitochondrial responses in rodent.

| HFD Duration | Mitochondrial Responses | Reference |
| --- | --- | --- |
| 2 weeks | Increased mRNA expression of mitochondrial dynamic genes, and fatty acid transport genes and uncoupling protein genes. Increased Drp1 protein content in skeletal muscles. | [61] |
| 15 days | Decreased activity of respiratory chain enzymes complex I, II, IV, and V in liver. Elevated GLUT1, 3 protein expression, reduced GLUT2, 4 protein expression in liver, skeletal and adipose tissue. | [62] |
| 3 weeks | Increased mitophagy in heart. Slightly lower mitochondrial enzyme activity, reduced mRNA for genes involved in OXPHOS and mitochondrial biogenesis in skeletal muscles. | [63], [64] |
| 4 weeks | Glucose intolerance, insulin resistance, increased mitochondrial biogenesis and impaired ADP sensitivity in skeletal muscles. | [65]–[67] |
| 8 weeks | Decreased mitochondrial density, impaired glucose metabolism in skeletal muscles. | [68] |
| 2 months | Reduced autophagy but continuously increased mitophagy in heart. | [63] |
| 10 weeks | Decreased maximal mitochondrial respiration, increased Fis1 level in skeletal muscles. | [69], [70] |
| 12 weeks | Reduced mitochondrial metabolic flexibility in skeletal muscles. | [71] |
| 16 weeks | Increased MAM formation, mitochondrial $\text{Ca}^{2+}$ overload, increased ROS generation, impaired insulin action and abnormal glucose metabolism in liver. Decreased mitochondrial membrane potential and ATP production, impaired glucose tolerance, increased Fis1 and Drp1 levels in skeletal muscles. | [51], [68], [70] |
| 28 weeks | Decreased mitochondrial energy production and biogenesis, changed mitochondrial morphology, decreased mRNA levels for mitochondrial dynamic genes, decreased MFN1, MFN2, OPA1 protein content, increased phosphorylated Drp1 and Fis1 protein levels in heart and heart hypertrophy. | [72] |
| 42 weeks | Abnormal metabolism, heart dysfunction, increased ER stress, decreased autophagy and mitophagy (decreased PINK1 and Mfn2) in heart. | [73] |

**Figure 1.**
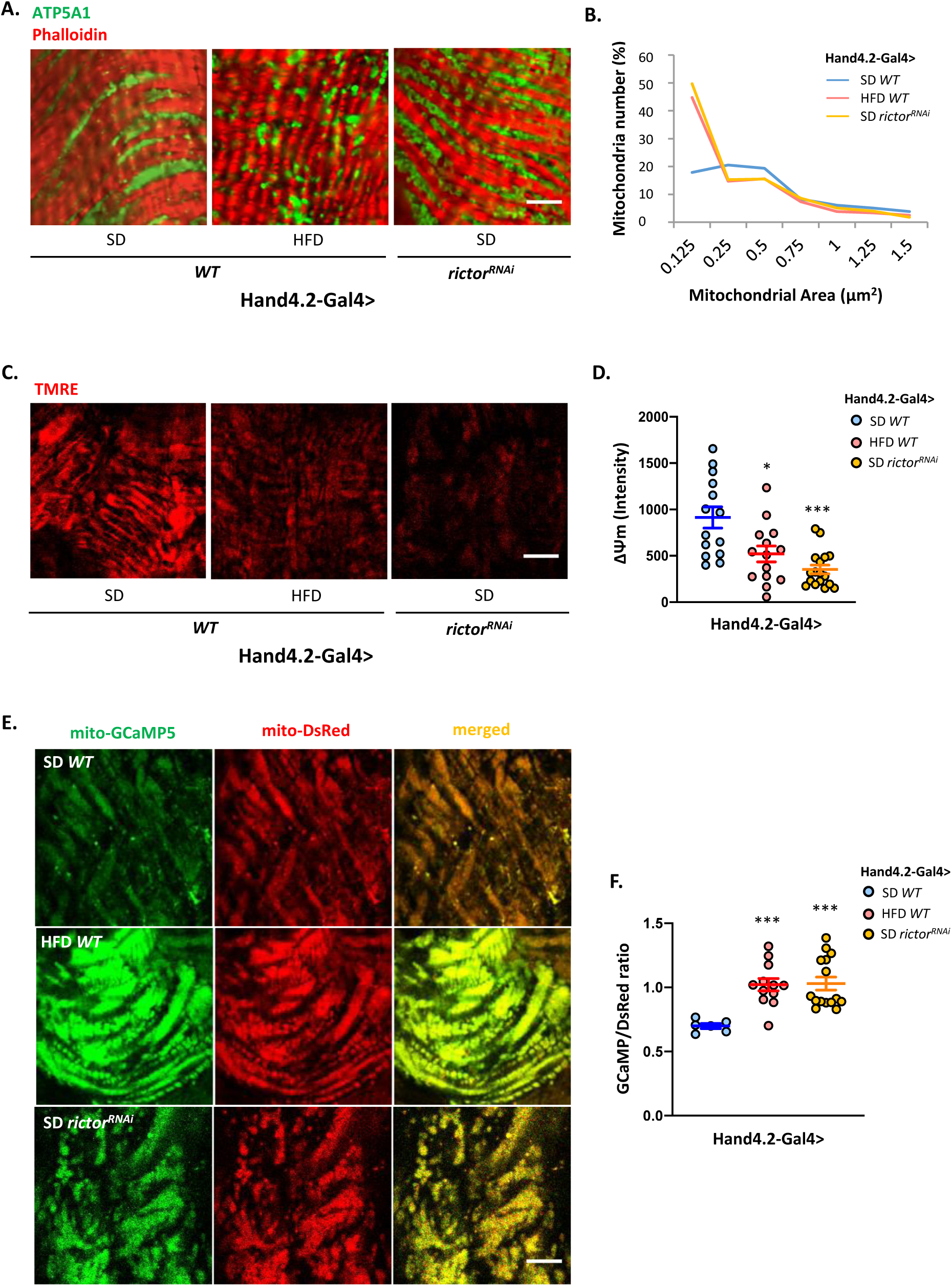
HFD and *rictor* knockdown alters mitochondrial physiology in *Drosophila* heart. (A) Area of mitochondria in the heart of wildtype or cardiac-specific *rictor* knockdown upon 5 days of HFD feeding. ATP5A1 antibodies were used to detect mitochondria and phalloidin was used to stain F-actin. (B) The group histogram data for mitochondrial area shown in (A). (C) Mitochondrial membrane potential measured by TMRE in the heart of wildtype or cardiac-specific *rictor* knockdown upon 5 days of HFD feeding. (D) The intensity profile of TMRE measured in (C). (E) Wildtype or *rictor* knockdown heart expressing mito-GCaMP5, and mito-DsRed upon 5 days of HFD feeding. (F) mito-GCaMP5 signal is normalized with mitochondrial mass (mito-DsRed) to represent the mitochondrial calcium level in heart. Flies were cultured at 40% relative humidity. *Hand-gal4* driver was used to drive gene expression specifically in cardiac tissues (cardiomyocytes and pericardial cells). Scale bar is 5 μm. N=4-6 and 3 ROIs were selected for each heart sample. Student t-test (* p<0.05, *** p<0.01).

Calcium (Ca^2+^) plays a critical role in regulating mitochondrial energy production and apoptosis. Besides, mitochondrial Ca^2+^ uptake has been closely linked to the regulation of mitochondrial dynamics and membrane potential. Mitochondrial membrane potential also serves as the driving force for Ca^2+^ uptake into the mitochondrial matrix. Thus, we next measured the mitochondrial Ca^2+^ levels in the HFD heart by normalizing mitochondrial-targeted Ca^2+^ reporter mito-GCaMP5 to mitochondrial mass (DsRed-mito). Our results indicated that HFD significantly increased the mitochondrial Ca^2+^ levels in the fly heart (Figure 1E, F).

Finally, we investigated how does HFD affects heart contractile function. In line with a previous study (Ryan T Birse et al., 2010), we also observed cardiac dysfunction reminiscent to a restrictive heart under HFD treatment [35]. Specifically, HFD slightly reduced diastolic diameter (DD) and significantly diminished diastolic interval (DI), and fractional shortening (FS) (Figure 2A-C) Collectively, HFD affects cardiac mitochondrial physiology as well as heart function in *Drosophila*.

**Figure 2.**
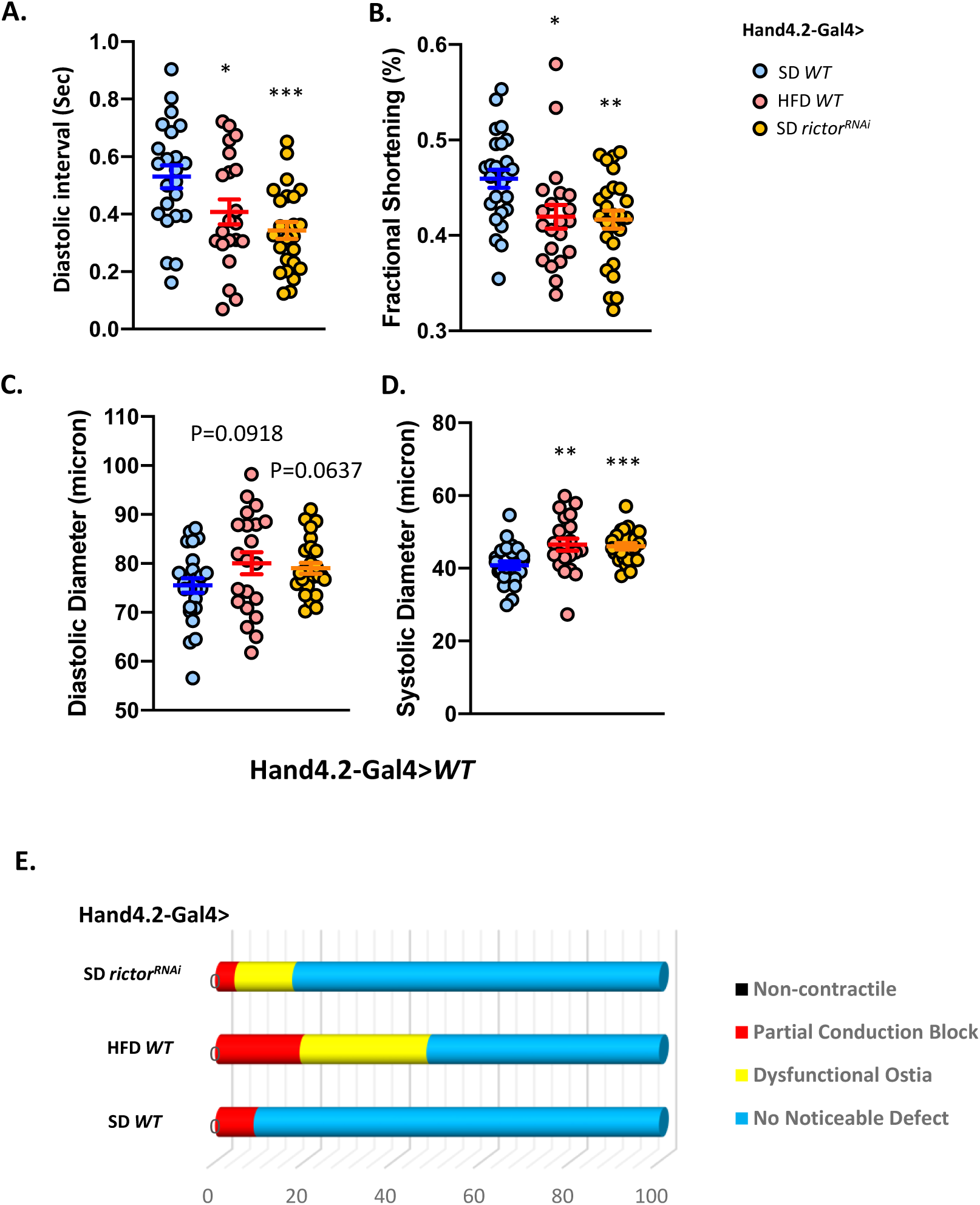
HFD alters cardiac function in *Drosophila*. (A) Diastolic interval (DI), (B) fractional shortening (FS), and (D) diastolic diameter (DD) of wildtype or *rictor* knockdown heart upon 5 days of HFD feeding. Flies were cultured at 40% relative humidity. *Hand-gal4* driver was used to drive gene expression specifically in cardiac tissues (cardiomyocytes and pericardial cells). N=21-25. Student t-test (* p<0.05, ** p<0.01, *** p<0.001, ns: not significant).

Previous studies suggest that MTORC2 plays a role in HFD-induced obesity and MTORC2/rictor deficiency displays HFD-related phenotypes (Bae et al., 2016; Chellappa et al., 2019; Cybulski et al., 2009; Dadalko et al., 2015; Mao & Zhang, 2018). Similarly, we also observed that cardiac-specific *rictor* knockdown resulted in changes similar to HFD-induced alterations of mitochondrial physiology and cardiac contractile patterns. We found that knocking down *rictor* in the heart induced mitochondrial fragmentation, dissipation of mitochondrial membrane potential, Ca^2+^ levels, and adversely affects cardiac function, including reducing DI, FS, and DD (Figure 2A-C). Consistently, we also found that the MTORC2/rictor protein level in the heart is reduced by five days of HFD treatment (Figure 3A, B). Therefore, reduced MTORC2/rictor activity in the heart at least partially takes part in regulating HFD-induced alterations in mitochondrial physiology and cardiac function in *Drosophila*.

**Figure 3.**
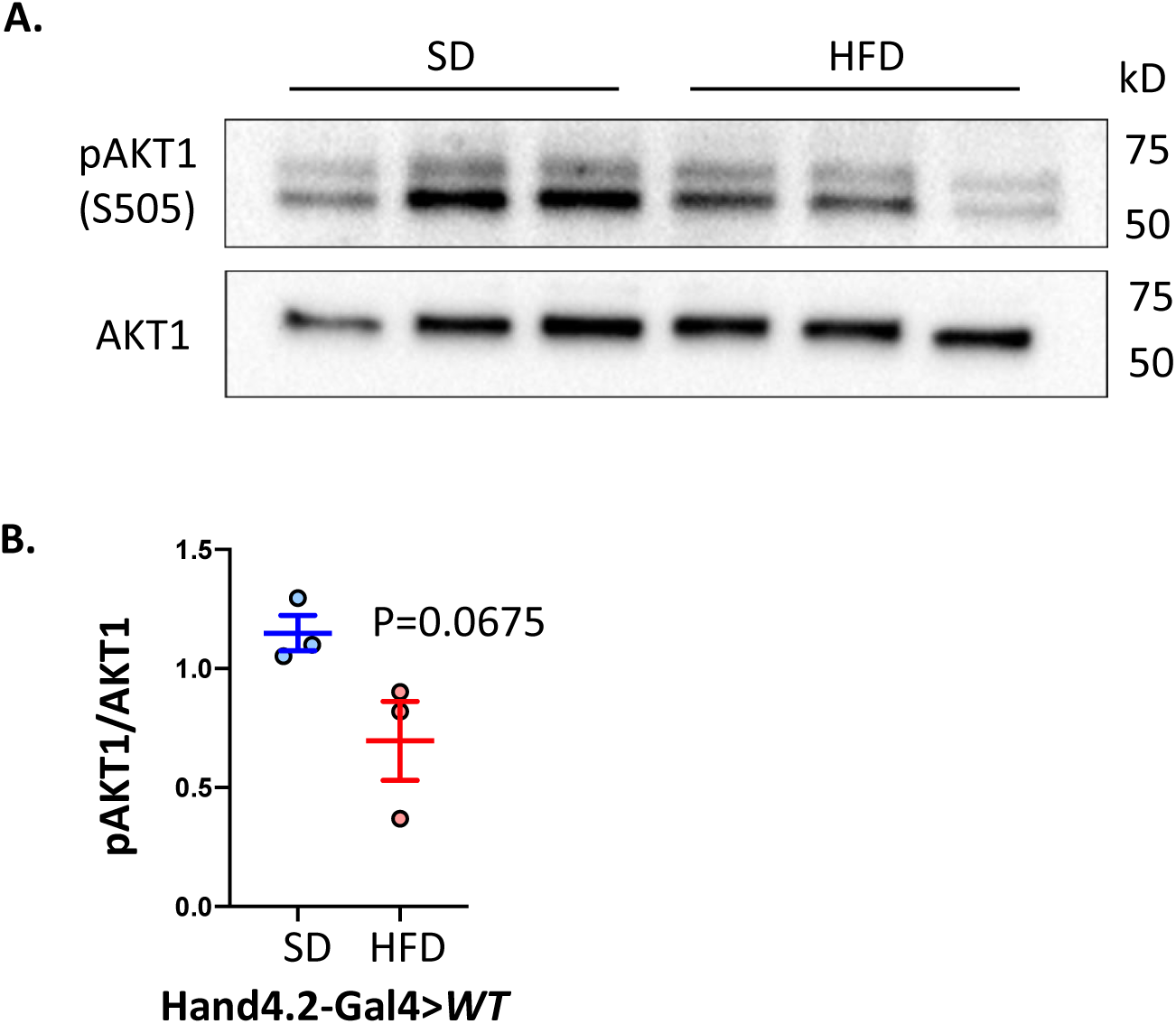
HFD reduces rictor activity in *Drosophila* heart. (A) Western blot analysis on Akt1 phosphorylation of hearts dissected from wildtype flies treated with 5 days of HFD. (B) The level of Akt1 phosphorylation is normalized to total Akt1 protein. Flies were cultured at 40% relative humidity. *Hand-gal4* driver was used to drive gene expression specifically in cardiac tissues (cardiomyocytes and pericardial cells). N=3 and 25-27 hearts were collected for each sample. Student t-test.

At last, we verified whether increasing MTORC2/rictor activity in the heart could rescue the HFD-induced mitochondrial and cardiac dysfunction. Our results suggested that cardiac-specific overexpression *rictor* prevents HFD-induced mitochondrial fragmentation (Figure 4A, C), dissipation of mitochondrial membrane potential (Figure 4B, D), and cardiac dysfunction (Figure 5A, B). Therefore, we concluded that MTORC2/rictor could provide cardio-protection in response to HFD.

**Figure 4.**
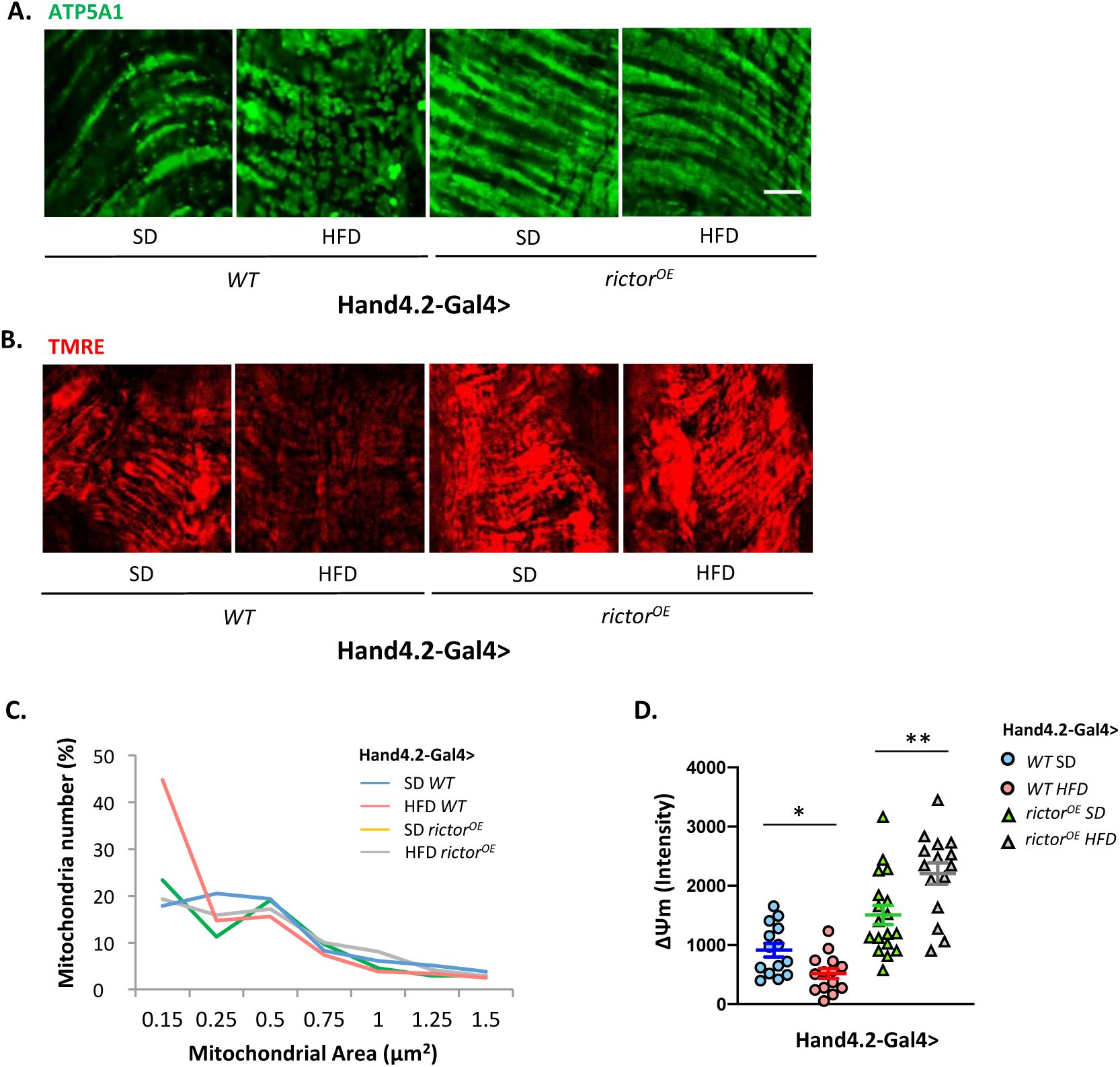
Overexpressing *rictor* rescues HFD-induced mitochondrial physiology alteration in *Drosophila* heart. (A) Mitochondrial size and (B) mitochondrial membrane potential in hearts of wildtype (*Hand4*.*2-Gal4>WT*) and cardiac-specific *rictor* overexpression flies upon 5 days SD or HFD feeding. ATP5A1 antibodies were used to detect mitochondria. (C) The group histogram data for mitochondrial area of (A). (D) The intensity profile of TMRE measured in (B). Flies were cultured at 40% relative humidity. *Hand-gal4* driver was used to drive gene expression specifically in cardiac tissues (cardiomyocytes and pericardial cells). Scale bar is 20 μm. N=5 and 3 ROIs were selected for each heart sample. Student t-test (* p<0.05, ** p<0.01).

**Figure 5.**
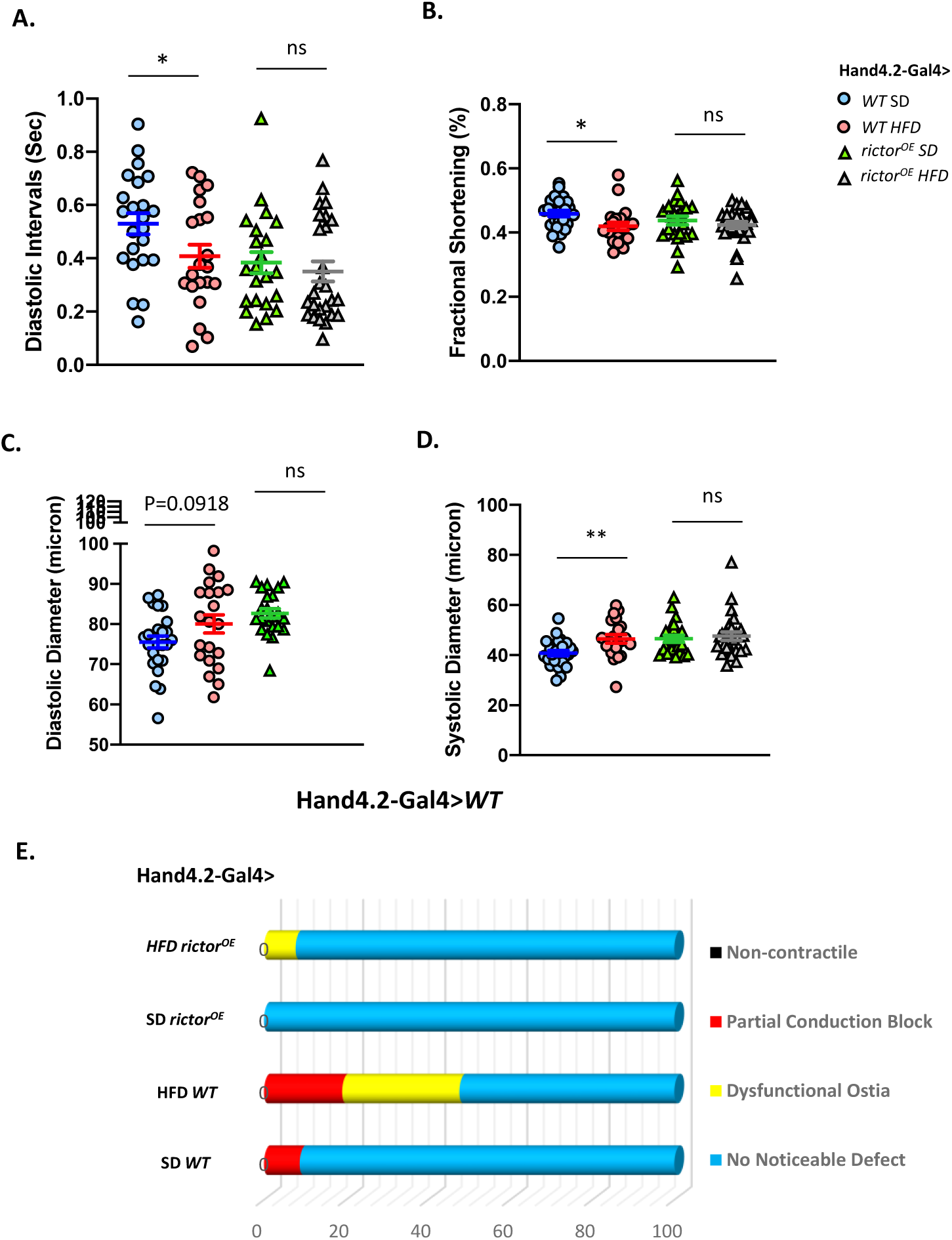
*rictor* overexpression rescues HFD-induced cardiac dysfunction in *Drosophila*. (A) Diastolic interval (DI) and (B) fractional shortening (FS) of wildtype or *rictor* overexpression heart upon 5 days of HFD or SD feeding. Flies were cultured at 40% relative humidity. *Hand-gal4* driver was used to drive gene expression specifically in cardiac tissues (cardiomyocytes and pericardial cells). N=21-25. Student t-test (* p<0.05, ns: not significant).

## Discussion

In this study, we evaluated the mitochondrial physiology and the contractile function in *Drosophila* heart when exposing to a HFD compared to a SD. Our results showed that HFD reduces mTORC2/rictor activity and induces mitochondrial fragmentation, dissipation of membrane potential, mitochondrial Ca^2+^ levels, and contractile dysfunction in Drosophila heart. Knocking down mTORC2/rictor in heart phenocopied above HFD-induced changes even on a SD, while overexpressing mTORC2/rictor in heart abolished above HFD-induced mitochondrial and cardiac dysfunction. Our study revealed a novel role of mTORC2/rictor in regulating cardiac mitochondrial physiology and HFD-induced cardiac dysfunction.

Mitochondria are known to change their architecture to meet the bioenergetic needs under different nutrient environments. Generally speaking, mitochondria tend to remain fragmented under lipid overload or other rich-nutrient environment and elongated under starvation conditions[36], [37]. This nutrient-induced mitochondrial fission supports “energy wasting” by enhancing uncoupling and basal protein conductance, which helps an increase in mitochondrial respiration meanwhile, a decrease in ATP synthesis efficiency[36], [37]. The mechanism by which mitochondrial fragmentation enhances uncoupling is not yet understood. One of the possibilities is that fragmentation might represent a change in cristae structures that allows the increased nutrient import and prevents mitochondrial ATP synthase dimerization[37], [38]. Even though mitochondrial fission seems to provide an adaptive response initially under a HFD, long-term exposure to nutrient overload such as a HFD still leads to increased ROS production, which is the major contributor to insulin resistance and mitochondrial dysfunction[39].

Mitochondrial membrane potential is highly correlated with the mitochondrial respiration rate[40]. The effect of nutrient overload on mitochondrial respiration and membrane potential remains controversial. Some studies suggest that nutrient excess increases mitochondrial respiration and membrane potential[41], [42], whereas others indicate that mitochondrial respiration is impaired by nutrient overload[17], [43]. A recent study in *Drosophila* skeletal muscles showed that mitochondrial respiration increases after two days on a HFD, followed by a significant decrease in mitochondrial respiration after four days of a HFD feeding. They demonstrated that the increased carbohydrates oxidation might contribute to the initial mitochondrial respiration increase since carbohydrates are the main fuel sustaining mitochondrial metabolism in muscles[44]. After continuous exposure to a HFD, the metabolic inflexibility occurs due to accumulated FFAs and depleted carbohydrates. Ultimately, the impairment of mitochondrial respiration ensues[17]. This evidence suggests that HFD treatment duration is one possible reason for the previous divergent results on mitochondrial membrane potential measurement. In addition, different cell types act distinctly to sense nutrients and utilize energy. For instance, nutrient utilization and its availability have greater control over mitochondria respiration and membrane potential in nutrient sensors such as beta cells. In contrast, ATP turnover significantly influences mitochondrial respiration and membrane potential in cells under high ATP demand such as muscle cells[37]. Therefore, the HFD-induced dissipation of cardiac mitochondrial membrane potential observed in our study is more likely to represent a deleterious condition caused by reduced ATP turnover, where the energy to support the contractile function is compromised. Unlike skeletal muscle, cardiomyocyte generates mostly ATP via fatty acid oxidation[6], which might explain why we did not see an increase in mitochondria membrane potential upon two days of HFD feeding.

Ca^2+^, a well-known regulator for mitochondrial function, is imported into the mitochondria matrix driven by membrane potential[40]. Once the Ca^2+^ enters the mitochondrial matrix, it controls the activities of several dehydrogenases in Kreb cycles, therefore regulates mitochondrial bioenergetics[45]. However, on the other hand, under the Ca^2+^ overload condition, increasing Ca^2+^ levels within the matrix could also promote mitochondrial permeability transition pore (mPTP) opening and dissipate mitochondria membrane potential[46]. Beyond mitochondrial energy metabolism, mitochondrial Ca^2+^ also promotes apoptosis under various stress conditions[47]. Recently, increasing evidence also suggests crosstalk between mitochondrial Ca^2+^ uptake and mitochondrial fission since both of these processes require the proximity between ER and mitochondria[48]–[50]. Collectively, mitochondrial Ca^2+^ is involved in a wide range of mitochondrial functions including mitochondrial fission and membrane potential. A recent study showed that obesity leads to increased mitochondrial Ca^2+^ uptake from ER via the MAM connections[51], and mitochondrial Ca^2+^ overload is known to lead to mPTP opening that triggers cardiac reperfusion injury[52], [53]. Indeed, HFD has shown to increase the vulnerability of hearts to ischemic reperfusion[54]. Therefore, the HFD-induced cardiac dysfunction observed in our studies might be due to deregulated mitochondrial Ca^2+^ uptake. Altogether, based on previous studies and our results, we speculate that the decreased mitochondrial function observed on five days of a HFD is not a regulated adaptive process but instead caused by damaging effects caused by nutrient excess.

To the best of our knowledge, there is only one study investigated the direct regulation between mTORC2 and mitochondria[26] despite some of the indirect evidence suggesting that mTORC2 is involved in mitochondrial quality control[55], [56]. In that specific study, they found that mTORC2 localizes to MAM to regulate its integrity and inhibits ER calcium release through IP3R. Therefore, testing whether mTORC2 provides cardioprotection via regulating MAM integrity and Ca^2+^ flux in response to a HFD could be one of the future directions. Additionally, our previous study has shown that mTORC2 slows cardiac aging through activating autophagy[57]; it is possible that mTORC2 also activates mitophagy to protect the heart from a HFD. In summary, our studies revealed a novel role of mTORC2 in the regulation of mitochondria in a HFD heart, study the mechanistic link between mTORC2 and mitochondria in the future could provide new insights for understanding mTORC2 and obesity-induced cardiovascular diseases.

## Materials and Methods

### Fly Husbandry and Stocks

Flies were maintained at 25°C, 60% relative humidity and 12 h light/dark. Female flies (1-2 weeks of age) were fed on a SD (agar-based diet with 0.8% cornmeal, 10% sugar, and 2.5% yeast) or a HFD (SD supplemented with 20% w/v coconut oil) for 5 days at constant densities (5 flies per vial). Fly stocks used in the present study are: *UAS-rictor RNAi* (BDSC, 31527), *Hand4*.*2-gal4* [58], *UAS-rictor* [59], *UAS-mito-GCaMP5/DsRed-mito* (Gift from Fumiko Kawasaki, Pennsylvania State University). *ywR* flies were used as control or wild-type (*WT*) flies.

### Fly Heartbeat Analysis

To measure cardiac function parameters, semi-intact *Drosophila* adult fly hearts were prepared according to previously described protocols [60]. In this study, we used the previous published fly heartbeat analysis [57]. In brief, flies were dissected to expose their hearts in oxygenated artificial hemolymph (AHL). Then high-speed digital movies of heartbeats were taken, and analyzed for DI, FS, etc.

### Immunostaining and Imaging

To investigate mitochondrial morphology in heart, we used ATP5A1 antibody (1:200; Invitrogen 15H4C4), which marks the mitochondrial ATP synthase. We used Alexa Fluor 594-conjugated phalloidin for F-actin staining (Thermo Fisher Scientific, A12381). All fluorescence-conjugated secondary antibodies were from Jackson ImmunoResearch (Alex Fluor 488).

For immunostaining, adult female flies were collected and dissected in AHL. Hearts were then incubated in relaxing buffer (AHL with 10mM EGTA) briefly to inhibit contractions. After fixing in 4% paraformaldehyde for 15 min at room temperature (RT), hearts were washed in PBS with 0.1% Triton X-100 (Fisher Scientific, BP151-100) (PBST) and then blocked in 5% normal donkey serum (NDS; Jackson ImmunoResearch, 005-000-121) diluted in PBST for 1 h at RT. Hearts were then washed with PBST and incubated overnight at 4°C with primary antibodies diluted in 5% NGS. After washing with PBST, the samples were incubated for 2 h at RT with appropriate fluorescence-conjugated secondary antibodies. Hearts were mounted in ProLong Diamond antifade reagent (Thermo Fisher Scientific, P36361) before being imaged using a FV3000 Confocal Laser Scanning Microscope (Olympus).

For image analysis and quantification, fluorescence images were analyzed in Olympus cellSens software. The mitochondria in a selected region of interest (ROI, ∼400 µm^2^) within heart tube were measured with the “Measure and Count” module in Olympus cellSens software. To quantify the mitochondria size, the area for each object/mitochondrion was measured and plotted in a distribution plot.

### TMRE staining

Flies were anesthetized and dissected in cold AHL. Hearts were then incubated in TMRE staining solution, consisting of 100nM of TMRE (Invitrogen, T668) in AHL for 12 min at RT. Samples were then rinsed twice for 30 s each wash with s solution consisting of 25 nM of TMRE in AHL. Hearts that attached to abdomen were quickly mounted in the same medium onto the slide and imaged within 15-20 min using identical setting on the confocal microscope. quantification of TMRE staining is done using cellSens, where mean intensity profile for the TMRE stains were quantified.

### Measurement of Mitochondrial Calcium

The Hand4.2-gal4 driver was used to drive the expression of UAS-mito-GCAMP5 and UAS-mito-DsRed reporter combination in adult heart. Flies were dissected to expose hearts in AHL, then the hearts that attached to the abdomen were immediately placed on the slides for live imaging. Images were taken with FV3000 Confocal Laser Scanning Microscope with a 100x oil-immersion objective lens. The mitochondrial calcium detected by mito-GCaMP5 were normalized with the UAS-mito-DsRed, which represents the mitochondrial mass.

### Western Blotting for MTORC2

The phosphorylation of AKT is used to represent MTORC2 activity [57]. 25-28 *Drosophila* adult hearts were collected for each sample. RIPA lysis buffer (Thermo Fisher Scientific, PI36978) was used to extract protein sample. Supernatants were collected and loaded onto Mini-PROTEAN precast gels (Bio-Rad Laboratories, 456–1095) using standard procedures. Blots were then incubated with primary and secondary antibodies. Primary antibodies used in this study included *Drosophila* p-Akt1 (Ser505) (1:1000) (CST, 4054) and AKT1 (Pan) (1:2000) (CST, 4691). All HRP-conjugated secondary antibodies are from Jackson ImmunoResearch. The blots were visualized with Pierce ECL Western Blotting Substrate (Thermo Fisher Scientific, PI34577). The images were analyzed by Image Lab.

### Statistical Analysis

GraphPad Prism (GraphPad Software) was used for statistical analysis. To compare the mean value of treatment groups versus that of control, either student t-test or one-way ANOVA was performed using Tukey multiple comparison. In SOHA analysis, the outliers were identified using Robust regression and Outlier removal (ROUT) method (Q = 1%) prior to the data analysis.

## Notes

### Competing Interest Statement

The authors have declared no competing interest.

